# Adapting the OpenFlexure Microscope for Affordable Live-Cell Imaging

**DOI:** 10.64898/2026.02.02.703252

**Authors:** Jodie R Malcolm, Olympia Physouni, Lola K Kelly, Juliana B T Carnielli, Sandy MacDonald, Mark Bentley, Stephen P Howarth, Stuart E Lacy, Alastair P Droop, Matthew Reilly, James P J Chong, Ana Paula C A Lima, Jeremy C. Mottram, Laura Wiggins, William J Brackenbury, Peter J O’Toole

## Abstract

Live-cell imaging (LCI) captures dynamic cellular behaviours missed in fixed samples. Researchers in resource-limited settings are underrepresented in the adoption of LCI due to the prohibitive cost of dedicated systems. We adapted the 3D-printed OpenFlexure Microscope (OFM) for LCI in humid tissue culture incubators. We relocated electronics to prevent corrosion, and printed in acrylonitrile styrene acrylate (ASA) to improve image stability in prolonged experiments. Current-limiting resistors reduced heat buildup inside sealed incubators, while our simple graphical user interface (GUI)-based application enabled rapid setup of timelapse experiments. Using a 48-hour docetaxel treatment of breast cancer cells, we validated how the LCI-OFM seamlessly fits into established bioimaging pipelines to generate biologically meaningful data. We further demonstrated broad applicability of the LCI-OFM by imaging *Leishmania*-infected macrophages in a containment level 3 laboratory and gut fungi in an anaerobic environment. This accessible platform expands opportunities for resource-limited researchers to study locally-relevant health challenges.

## Introduction

Timelapse microscopy is essential for studying dynamic biological processes. Many commercial high-end imaging platforms exist that enable continuous, label-free monitoring of living cells. These systems reveal important phenotypic transitions that are overlooked or missed when studied in fixed-sample workflows, including subtle changes in cell morphology, motility, proliferation or response to drug treatment ^1–7^. For many researchers, datasets monitoring cells over time provide a superior understanding of their behaviour, and when these data are integrated with advanced computational tools including machine learning-based phenotyping pipelines, quantitative insights beyond the capabilities of manual inspection alone can be extracted ^8^.

Despite the scientific value of live-cell imaging (LCI), access to dedicated systems remains highly unequal. Researchers in resource-limited settings, such as in low-to middle-income countries (LMICs), are often excluded by the prohibitive cost of these platforms, restricting their ability to generate high-resolution time-series data necessary to advance research addressing locally-relevant health concerns ^9,10^. This disparity has motivated the wider microscopy community to develop frugal microscopy solutions to democratise access to advanced imaging ^11^. At the extreme, Foldscope is an ultra-low-cost origami-based microscope with brightfield, darkfield and fluorescence capabilities that has been brought to fruition to enable large-scale manufacturing of frugal microscopes ^12^. CellScope combines a collection of optics, LED illumination and bluetooth controlled motors with 5 MP cameraphone devices to create a device capable of collecting images suitable for remote diagnoses of oral cancers ^13,14^. Glowscope can transform smartphones or tablets to functional fluorescence microscopes using re-purposed LED flashlights and lighting filters, and is capable of 10 μm resolution ^15^. Initiatives that repurpose smartphones into functional microscopes represent an accessible solution that broadens opportunities for research and remote diagnostics. Modular microscopes such as the OpenFrame and squid further provide researchers with an opportunity to configure custom imaging systems tailored to specific scientific questions with possibilities to upgrade or expand the capabilities of the microscope over the lifetime of the instrument ^16,17^. Collectively, these initiatives reflect a broader commitment to making microscopy more accessible and adaptable, and broadening global participation. The OpenFlexure Microscope (OFM) ^18,19^ has also been successful in breaking down these barriers and in particular stands out for being a highly customisable 3D-printed instrument, costing only a few hundred pounds. Despite the OFM being a valuable tool in medical settings ^19,20^, we identified limitations that prevent its use in long-term live-cell experiments, particularly those that will need to be carried out in routine humidified and temperature-controlled incubators to maintain the health of living cells, creating a gap between the promise of low-cost open-source microscopy and the practical requirements of robust, long-term LCI.

Here we present the LCI-OFM, achieved by making essential modifications to the original microscope hardware to improve environmental resilience and usability for timelapse experiments. Adaptations include isolating critical electronics from the incubator, protecting internal circuitry with a sealed chassis, incorporating current-limiting resistors, and identifying a durable polymer to print the main microscope chassis. We also created a simple timelapse-specific graphical user interface (GUI)-based application to enable intuitive control of stage movement and illumination.

We validated the usability of the redesigned system through a 48-hour treatment response of MDA-MB-231 breast cancer cells with the chemotherapeutic docetaxel, generating data compatible with established bioimage analysis workflows including Trackmate ^21^, cellpose ^22^ and the CellPhe toolkit ^8,23^. Additionally, we successfully operated the LCI-OFM in a containment level 3 laboratory to image *Leishmania*-infected macrophages, and within an anaerobic workstation to capture anaerobic gut fungi of the monocentric *Neocallimastix* genus, highlighting the microscope’s broad applicability across diverse biological environments.

By bridging the gap between open-source microscopy and the demands of modern LCI, our work provides an accessible tool for researchers in resource-limited settings to generate preliminary timelapse imaging data of sufficient quality to be segmented using CellPose, tracked using Trackmate, analysed using CellPhe, pursuing locally relevant biological questions, and strengthening access to future funding and imaging infrastructure. This offers a real opportunity for the development of live-cell imaging skills, preliminary data sets to access more established facilities, and to democratise the availability of live-cell imaging for all, and not just well funded labs.

## Results

### Essential modifications to the OpenFlexure hardware and electronic components to enable live-cell imaging in humid tissue culture incubators

The high resolution motorised OFM (V7) comprises three main parts (Figure 1a): (i) the base housing a Raspberry Pi 4 Model B and Sangaboard (V5); (ii) the main body which includes the flexure-based chassis, sample stage, *x*, *y* and *z* stepper motors and a Royal Microscopical Society (RMS)-thread Plan-corrected 40X 0.65 numerical aperture (NA) objective connected to a Raspberry Pi Camera Module (V2) and; (iii) the illumination fitting with a condenser lens and LED.

**Figure 1.**
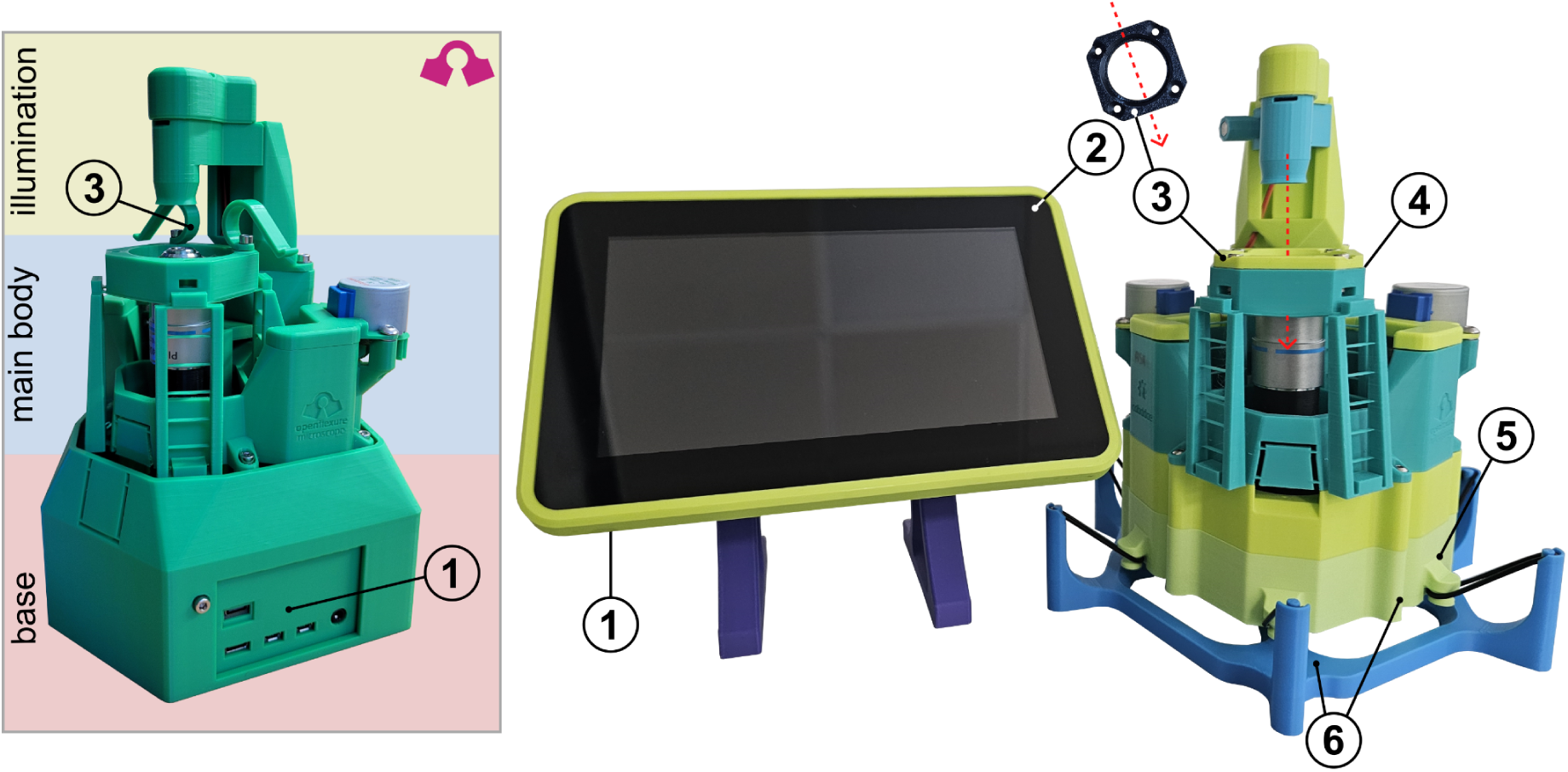
Overview of hardware modifications to the OpenFlexure Microscope to enable live-cell imaging. (a) The original design of the OFM can be compartmentalised into the base unit, the main body and the illumination fitting. (b) To enable live-cell imaging we redesigned the OFM to create a designated LCI-OFM: (1) The Raspberry Pi 4 model B and Sangaboard V5 have been relocated from the OFM mainbody to a custom-built case designed to also hold a (2) Raspberry Pi 2 Display. (3) Slide clips on the original OFM have been replaced with a stage-adaptation to support standard 35 mm culture dishes necessary for maintaining optimal conditions for living cells. (4) To account for increased height dimensions from the sampling vessel, a removable spacer has been added that raises the condenser and LED by 12 mm. (5) A new base has been designed to seal off the camera and motor connections from a humid incubator. (6) A hollow hammock support was designed to raise the microscope off the incubator shelf using O-rings to reduce vibration artefacts.

To enable LCI, we implemented adaptations essential for maintaining cell health during timelapse experiments (Figure 1b; Supplementary File 1). To support cell growth and maintain adequate volume of culture medium, we omitted the slide clips typically used for imaging fixed samples, and designed a printable insert that supports 35 mm culture dishes. The underside of the insert is recessed to maintain the distance between the objective and the imaging surface from the original configuration. To raise the height of the condenser arm and LED to fit taller culture vessels, we designed a 12 mm spacer that can be fixed between the illumination dovetail and the stage. As the new stage insert and illumination spacer are printed independently of the rest of the microscope, users have the option to interchange between slide clips and the dish holder depending on experimental requirements.

Placing the OFM inside a tissue culture incubator risks exposing electronic components to humid environments, which could cause corrosion or short-circuiting, resulting in complete failure of the instrument. To overcome this, we isolated the Raspberry Pi 4 Model B and Sangaboard from the main body of the microscope and designed a dedicated housing that sits outside of an incubator. This enclosure incorporates a Raspberry Pi 2 Display to enable easy control and real-time monitoring of experiments, and also removes the need for external monitors and HDMI connections making the setup of the OFM more streamlined. To preserve the Raspberry Pi Camera, ribbon and cables, we closed off the top and underside of the microscope with a new 3D-printable base, creating a physical barrier between fragile electronics and the harsh incubator environment. Additionally, by separating the Raspberry Pi from the optics module, any liquid spills would only require the 3D-printed shell to be replaced, while the lens can be sterilised and the Pi remains protected, making the design highly desirable for high-risk category 3 research.

Tissue culture incubators experience vibrations due to internal fans that are used to maintain uniform environmental conditions, or users opening and closing the incubator doors. Vibrations affect image quality, resulting in vibration-induced artefacts including blurring, image smearing, or frame drifting and shunting. To provide a cost-effective solution to vibration issues in timelapse imaging, we designed a hammock system that suspends the OFM above the incubator shelf using O-rings to provide passive vibration isolation. To ensure reliable operation in settings with unstable electricity provision, we tested the LCI-OFM using a powerbank as an uninterrupted power source, and confirmed that it runs seamlessly from an Anker 537 (PowerCore 24K) even when mains power is lost.

### A simple graphical user interface for timelapse setup

Operating the original OFM uses a custom Raspberry Pi operating system with OpenFlexureConnect, which provides the live preview, image capture and access to web-based control interfaces such as OpenFlexure Blockly for scripting and timelapse setup. While these interfaces support flexible, advanced control they are not optimised for rapid configuration of routine timelapse experiments. To streamline timelapse experiment workflows, we developed a user-friendly, LCI-focused GUI (LCI-GUI) that can be installed locally onto Raspberry Pi operating systems and enables direct camera access, rapid timelapse setup and continuous monitoring of experimental progress (Figure 2). Installing our LCI-GUI does not restrict terminal access to the Pi Camera, allowing users to retain complete control of the camera for more advanced or specialist microscope applications. Additionally, our LCI-GUI does not require an internet connection for use, and modifications to the LCI-GUI program can be installed offline, providing further customisation for the user.

**Figure 2.**
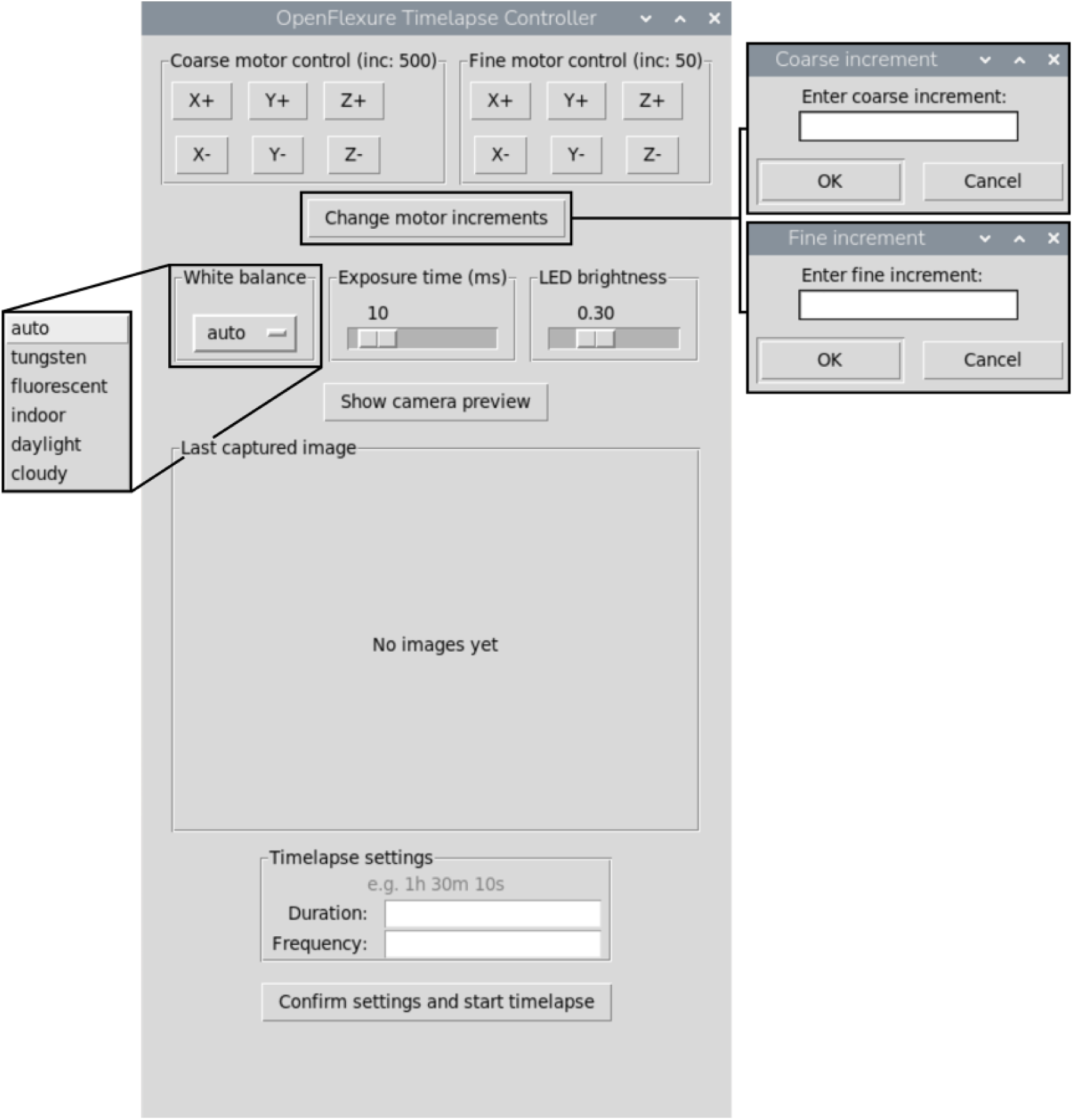
Simple graphical user interface designed for rapid timelapse set up. The LCI-GUI is designed to allow the user to operate the microscope locally to set up a timelapse experiment. Coarse and fine motor control allows stage positioning along the *x*, *y* and *z* axis with the option to change default increment size for each motor. White balance is pre-set to “auto”, but a drop-down menu allows the user to optimise the white balance setting based on the experimental requirements or preference. Similarly, exposure time (ms) and LED brightness can be adjusted using the relevant sliders for optimal image acquisition. “Show camera preview” opens a new window to allow the user to move around the sample to find a region of interest. Once the user is ready to start their timelapse, they can simply type in the duration (h:m:s) and the interval time (h:m:s) and then “confirm settings and start timelapse”. Throughout the timelapse, the last image to be acquired is shown in the “last captured image” box, enabling users to monitor the progress of their experiment in real time.

When the application opens, users are able to identify and focus a region of interest (ROI) using course and fine *x*, *y* and *z* motor controls (Figure 2). Users can also specify the size of motor increments, alter the white balance, and control exposure time and LED brightness. A live preview is displayed in a new window prior to starting a timelapse experiment. Timelapse scheduling is controlled by a user-specified duration and frequency, both in hours, minutes, and seconds. When a user has confirmed settings and started the timelapse, the “last captured image” box will update at the specified frequency to show the last picture taken during the experiment. Each timelapse is stored within its own timestamped directory containing every acquired image encoded as JPEG, which are also timestamped (e.g. “2024-12-19_15_25_26.jpeg).

Because the LCI-GUI lacks built-in colour-channel correction, raw timelapse images show uneven colour balance and vignetting. To address this, we generated a simple ImageJ macro (Supplementary File 2) that converts RGB images to 8-bit greyscale, producing uniform shading without need for calibration. As demonstrated using MCF-7 and MDA-MB-231 cells (Supplementary Figure 1), this correction improves image quality and consistency across the field of view. The macro also includes 50% down-scaling, substantially reducing file size while retaining important biological features. Smaller greyscale images lower storage demands and computational requirements, an important consideration for LMIC-feasible bioimaging workflows.

### Printing in acrylic styrene acrylonitrile is recommended for timelapse stability and durability of the microscope in humid tissue culture incubators

Initial experiments with the LCI-OFM experienced focus loss and optical drift away from specified ROIs (Supplementary Figure 2; Supplementary Video 1). We hypothesised that movement seen during these experiments could be explained by the chemical properties of the Polylactic Acid (PLA) used to print the microscope body. To minimise focal drift and to enhance the overall durability of the microscope, we explored the mechanical characteristics of PLA, PLA tough (PLA+), Acrylic Styrene Acrylonitrile (ASA) and Flame-Retardant Polycarbonate (PC-FR). Full details of the tensile strength, bending strength and Vicat Softening Temperature (VST) are described in Supplementary Table 1.

To assess which polymer would provide superior microscope durability, we printed the LCI-OFM main body using each material and used the LCI-GUI to initiate a 24-hour timelapse to monitor ROI stability. The first image captured at the start of the timelapse highlights the high resolution attainable using 3D-printed microscopes, with cell morphology, texture and nucleus clearly visible (Figure 3). Throughout the 24-hour timelapse, PLA and PLA+ both resulted in substantial focal drift, suggesting polylactic acid polymers are not suitable for use in humid tissue culture incubators at 37°C (Figure 3, Supplementary Video 2 and 3). PC-FR outperformed PLA and PLA+ with its ability to maintain positioning during the experiment (Figure 3). However, cells appeared to be “shaking” with each frame, as this polymer was strongly susceptible to vibrations from the incubator (Supplementary Video 4). These artefacts would be difficult to account for in bioimage analysis pipelines. Printing with ASA was superior compared to the other polymers, as there were fewer vibration artefacts and minimal focal drift during the experiment (Figure 3; Supplementary Video 5). Other polymers not explored, including PET may also perform well. However, the higher cost of PET makes it less suitable for resource limited settings, and the lower cost PETG is more difficult to handle in many printers. Therefore, we proceeded with ASA as the printing material for the LCI-OFM main frame.

**Figure 3.**
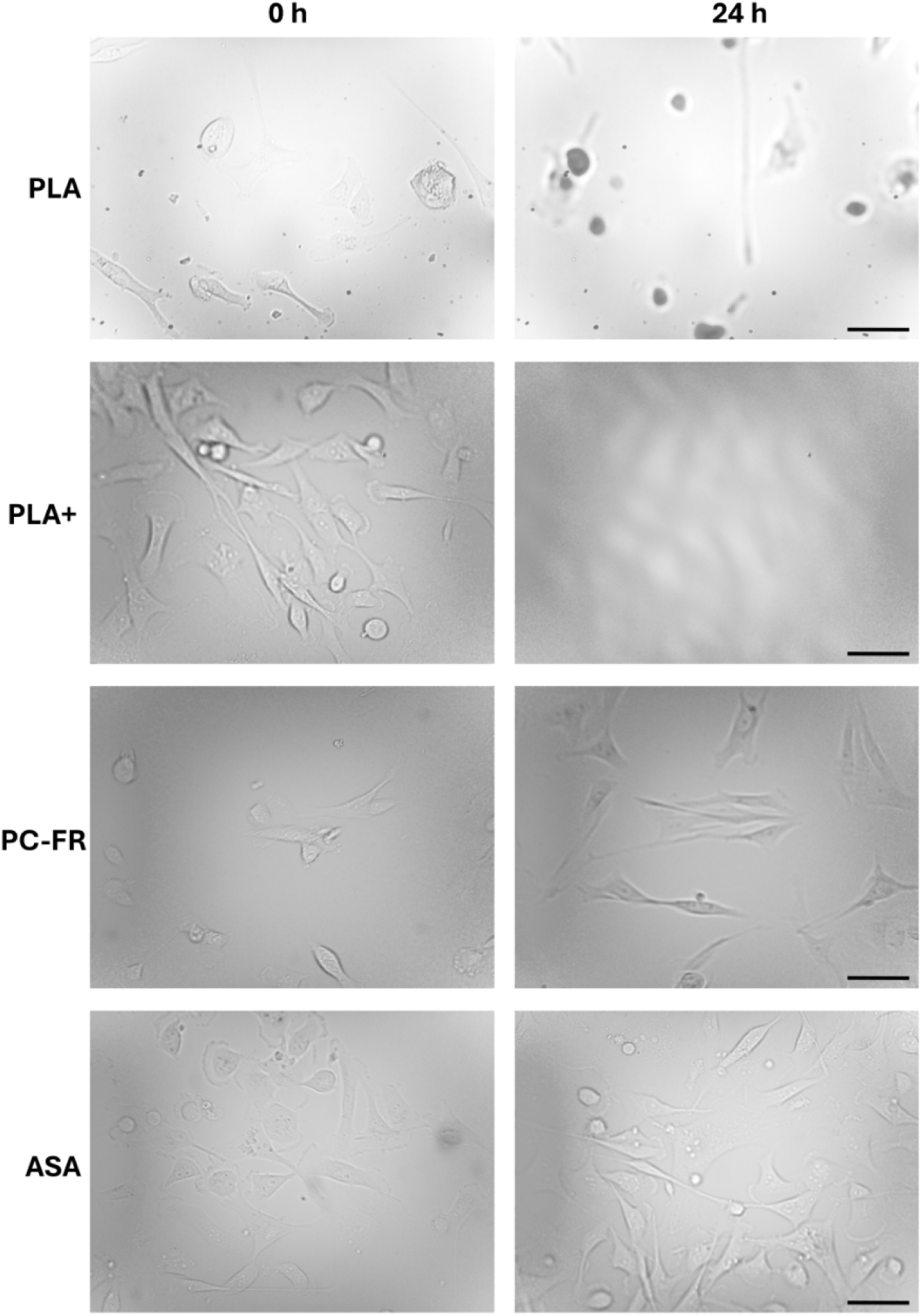
Testing common 3D-printing polymer materials for image quality during short timelapse experiments. MDA-MB-231 breast cancer cells were imaged every 6 min for 24 h at 40X magnification, using LCI-OFMs where the main frame was printed in either PLA, PLA+, PC-FR or ASA. The first and last image that was taken during a 24 h window is shown, to enable visual comparison of image quality and focal drift. Raw images were processed at the end of the timelapse using our ImageJ macro described above. Microscopes were placed in the tissue culture incubator at least 24 h before imaging to allow the metal and plastic components to equilibrate to the higher temperature and stabilise. Scale bar = 50 µm.

### Custom resistor plugins are essential for reducing stepper motor temperatures during prolonged timelapse experiments

Early timelapses found that the *x*, *y* and *z* stepper motors generated substantial heat while inside incubators, which could compromise cell health by disrupting the tightly controlled environment. To address motor heating, we designed and tested custom plugins with 15 ohm (Ω), 22 Ω or 27 Ω resistors to limit the current supplied - as heat dissipation is directly proportional to the square of the current ^24^ - while still delivering sufficient power for motor function (Figure 4a).

**Figure 4.**
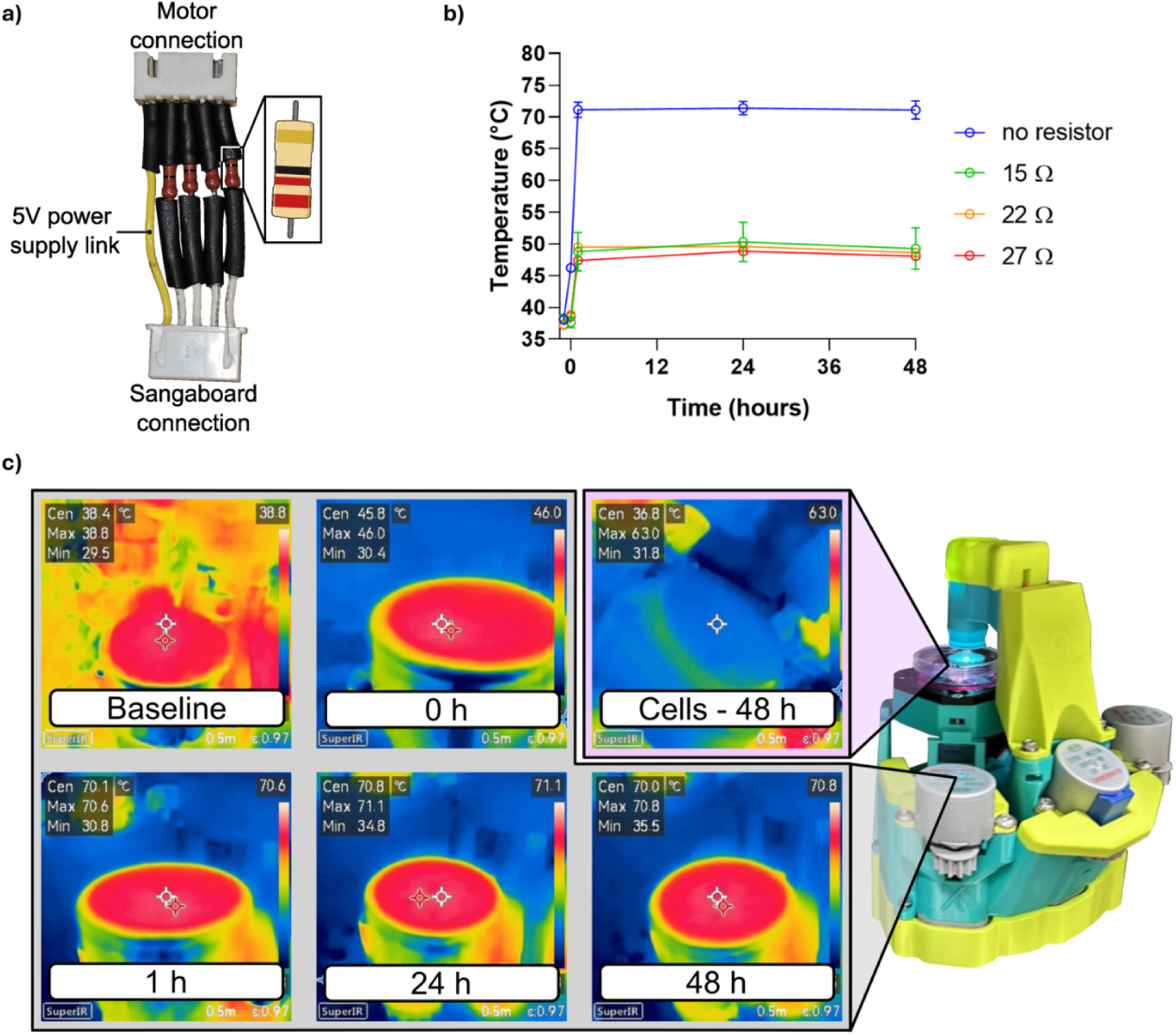
Controlling temperature of OFM stepper motors with resistors. a) A diagram showing the current limiting plugins. The image represents a plugin composed of four 22 Ω resistors and a 5 V power supply link wire, which is socketed into the *x*, *y* or *z* motor housing of the Sangaboard and connects to each individual motor extension. b) Temperature recordings of *x*, *y* and *z* motors powered without any plugins, or with plugins made of four 15 Ω, 22 Ω or 27 Ω resistors as shown in (a). Baseline readings were recorded after the modified OFM had been placed into a humid tissue culture incubator for 24 hours, before connecting the power supply. The 0 h measurement was taken immediately after the modified microscope had been powered on and a timelapse initiated (i.e., after moving around the sample with *x* and *y* axis motor movements to find a ROI, and adjusting focus with *z* axis motor). Subsequent temperatures of each motor were taken 1, 24 and 48 h post timelapse initiation. Recordings were taken with a handheld superIR resolution camera. Error bars are ± SEM. c) Representative images of thermal images captured using the Hikmicro Eco handheld superIR resolution camera. Displayed are the IR pictures taken of the *x* motor in a modified OFM that did not have any resistors, and of the cells from the same microscope at the end of a 48 hour timelapse experiment.

Without plugins, the power consumption per winding of the stepper motor was calculated to be 0.68 W. At the onset of the timelapse, the temperature of the motors increased from a baseline of 38.1 °C (± 0.5 SD) to 46.2 °C (± 0.8 SD) (Figure 4b, 4c). One hour after the timelapse was started, the temperature of the motors stabilised at 71.1 °C (± 2.1 SD), which was maintained at the 24 hour (71.4 °C, ± 1.8 SD) and 48 hour (71.1 °C, ± 2.5 SD) timepoints. Despite the high temperature of the stepper motors during the experiment, the vessel containing MDA-MB-231 cells remained an optimal 36.8 °C and the cells appeared to be healthy throughout the experiment (Supplementary Figure 2, Supplementary Video 6) evidencing that there were no negative localised effects. Adding four 15 Ω resistors into the plugins reduces motor current by 40%, and 0.24 W per energised winding. The baseline temperature of the motors with this lowest amount of resistance was 38.3°C (± 0.5 SD), and 37.6 °C (± 1.5 SD) immediately after timelapse initiation. The motors with the 15 Ω plugins stabilised after one hour at 48.8°C (± 5.3 SD), reaching 50.3 °C (± 5.4 SD) after 24 hours and settling at 49.3 °C (± 5.66 SD) following 48 hours. The power consumption for plugins with 22 Ω resistors was calculated to be 0.17 W per energised winding. The temperature decrease with 22 Ω plugins was marginally improved overall compared to the 15 Ω plugins, starting with a baseline temperature of 37.3 °C (± 0.4 SD) which increased to 38.9 7°C (± 0.2 SD) at the start of the timelapse. After one hour, the stepper motors were 49.5 °C (± 0.8 SD) which was sustained at the 24 hour (49.6 °C, ± 0.8 SD) and 48 hour (48.6 °C, ±0.9 SD) timepoints. For the plugins with 27 Ω resistors, power consumption per energised winding was decreased further to 0.14 W. Although these 27 Ω plugins were comparable to the 22 Ω plugins in terms of heat generation (Figure 4b), the motors were noticeably less responsive to commands from the LCI-GUI motor controls, suggesting that this setup limited power delivery to the stepper motors below the amount required for reliable use. Therefore, we opted to use the 22 Ω plugins in the final design of the LCI-OFM due to the improved heat reduction at the stepper motors without compromised functionality.

### The modified OpenFlexure is suitable for studying changes in cellular phenomics over time

We wanted to demonstrate that data obtained from the LCI-OFM are biologically relevant, and sufficient for seamlessly fitting into established bioimage analysis pipelines. We used our LCI-GUI to rapidly set up a 48-h timelapse of MDA-MB-231 breast cancer cells following treatment with either a vehicle control (0.01% DMSO) or 1 μM docetaxel, with a 6-min imaging interval. We observed that MDA-MB-231 cells treated with the vehicle maintained their characteristic elongated morphology and exhibited active migration and proliferation throughout the experiment, reflecting normal cytoskeletal dynamics and cell cycle progression (Figure 5a; Supplementary Video 7) ^25^. Docetaxel-treated cells rapidly adopted a rounded, enlarged morphology consistent with microtubule stabilisation and mitotic arrest. These cells regularly failed cytokinesis and showed overall reduced motility, highlighting the cytostatic and cytotoxic effects of docetaxel on this breast cancer cell line (Figure 5a; Supplementary Video 8) ^26,27^. Importantly, these biologically-relevant dynamic characteristics were able to be captured using label-free imaging on our low-cost 3D-printed LCI-OFM.

**Figure 5.**
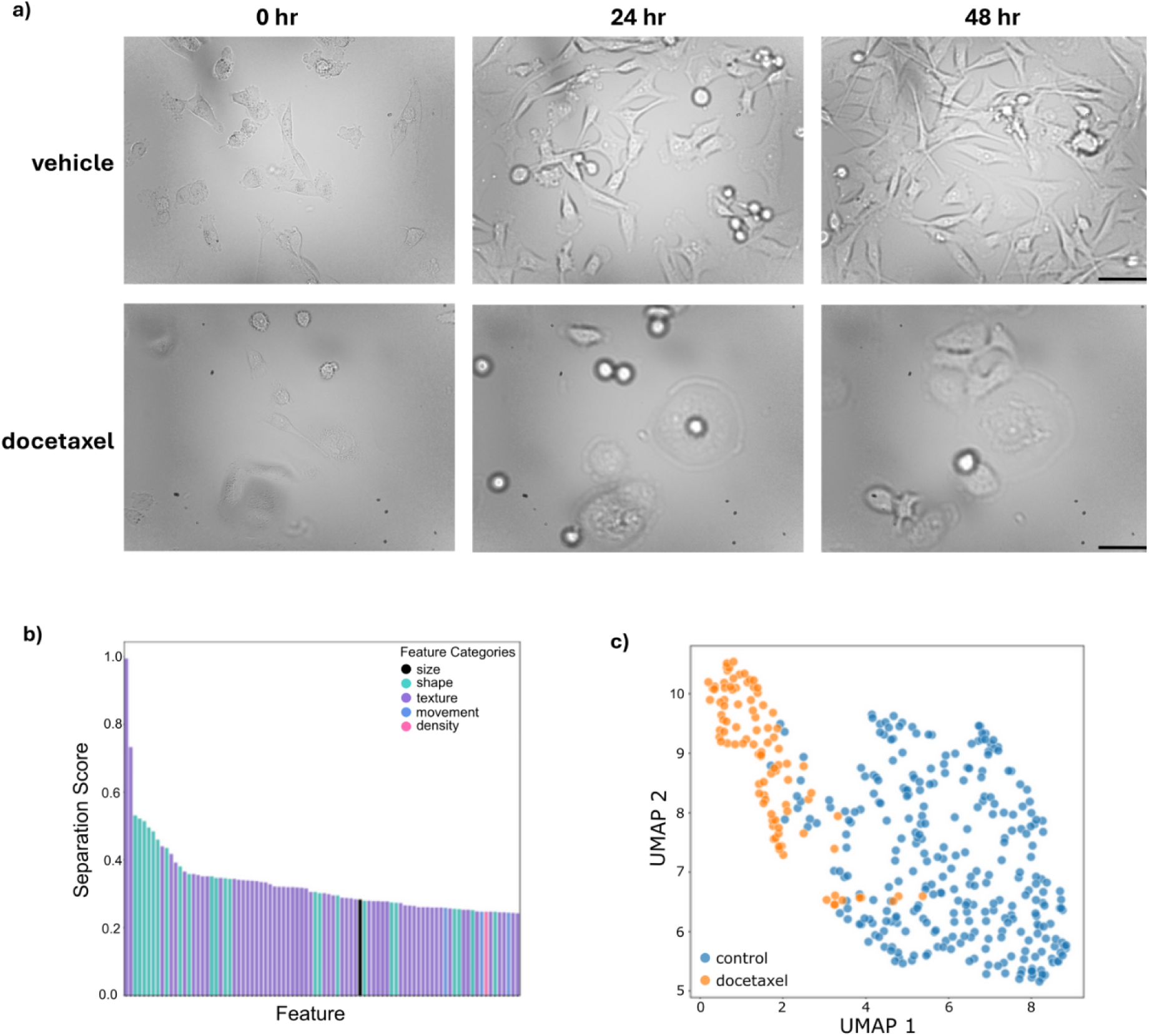
Validating the modified OFM as a cost-effective tool for live-cell imaging. (a) MDA-MB-231 breast cancer cells were treated with either 0.01% DMSO (vehicle) or 1 μM docetaxel and imaged at 40X magnification using the LCI-OFM. Images were taken every 6 min for 48 h and the experiment was repeated four times (N = 4). Raw images were processed at the end of the timelapse using the imageJ macro described above. LCI-OFMs were placed in the incubator 24 h before the start of the first timelapse experiment was initiated, and remained in place until completion of the fourth and final experiment (17 days total). Scale bar = 50 µm. Using a custom Cellpose segmentation model and Trackmate, we were able to utilise the CellPhe toolkit to extract features that are most discriminating between vehicle and docetaxel-treated MDA-MB-231 cells that had been imaged for 48 h using our LCI-OFM. (b) Separation scores and feature categories of the most discriminating features between the control and drug-treated breast cancer cells. (c) UMAP showing feature separation between vehicle (blue) and docetaxel-treated MDA-MB-231 cells (N = 4).

We next subjected these datasets to an established bioimage analysis pipeline. Previously we have developed CellPhe, a machine-learning-based pattern recognition toolkit that reliably identifies features pertaining to cell shape, size, texture, movement and density that best discriminate between populations in time-series experiments ^8,23^. Here, we used a custom Cellpose model ^22^ trained on MDA-MB-231 LCI-OFM timelapse experiments, TrackMate ^21^ and the CellPhe Toolkit to analyse four biological replicates of the MDA-MB-231 drug treatment assay. Our custom Cellpose model performed well at segmenting MDA-MB-231 cells with varied morphology in response to treatment (Supplementary Figure 3). Using the CellPhe toolkit, we identified 88 of 1061 features that discriminated between vehicle and docetaxel-treated cells. Texture and shape-related metrics dominated the discriminating features, whereas movement and density-based metrics contributed less (Figure 5b). These data align with the discriminating features previously identified between vehicle-and docetaxel-treated MDA-MB-231 cells using the high-end Phasefocus Livecyte instrument, where texture, shape and size metrics were also the most informative features between these two groups ^8^. Importantly, although the timelapse was recorded using a frugal 3D-printed microscope, the image quality was sufficient to clearly separate control and drug-treated breast cancer cells, underscoring that low cost instruments can yield biologically relevant data (Figure 5c).

### Broad Applicability of the LCI-OFM Across Diverse Biological Systems

Finally, we demonstrate that the LCI-OFM is a versatile imaging platform capable of operating across diverse biological systems, extending well beyond timelapse imaging of mammalian cells in standard tissue culture incubators.

Imaging in a containment level 3 laboratory is inherently challenging due to strict containment requirements, limited access, and restrictions on moving high-hazard pathogens to external facilities. These constraints limit the use of conventional imaging systems and necessitate workflows that minimise handling to maintain biosafety. By positioning the LCI-OFM directly inside category 3 incubators, we eliminate the need to transport hazardous samples and enable immediate imaging under physiologically relevant conditions. Using this approach, we visualised host-pathogen interactions in bone marrow-derived macrophages infected with late-logarithmic phase *Leishmania donovani* promastigotes. After 6 hours, non-internalised parasites were washed away and cultures were returned to the incubator for immediate imaging. The LCI-OFM clearly resolved early parasite uptake, showing the visible extracellular flagellum of a promastigote during macrophage entry (Figure 6a), capturing initial stages of the infection.

**Figure 6.**
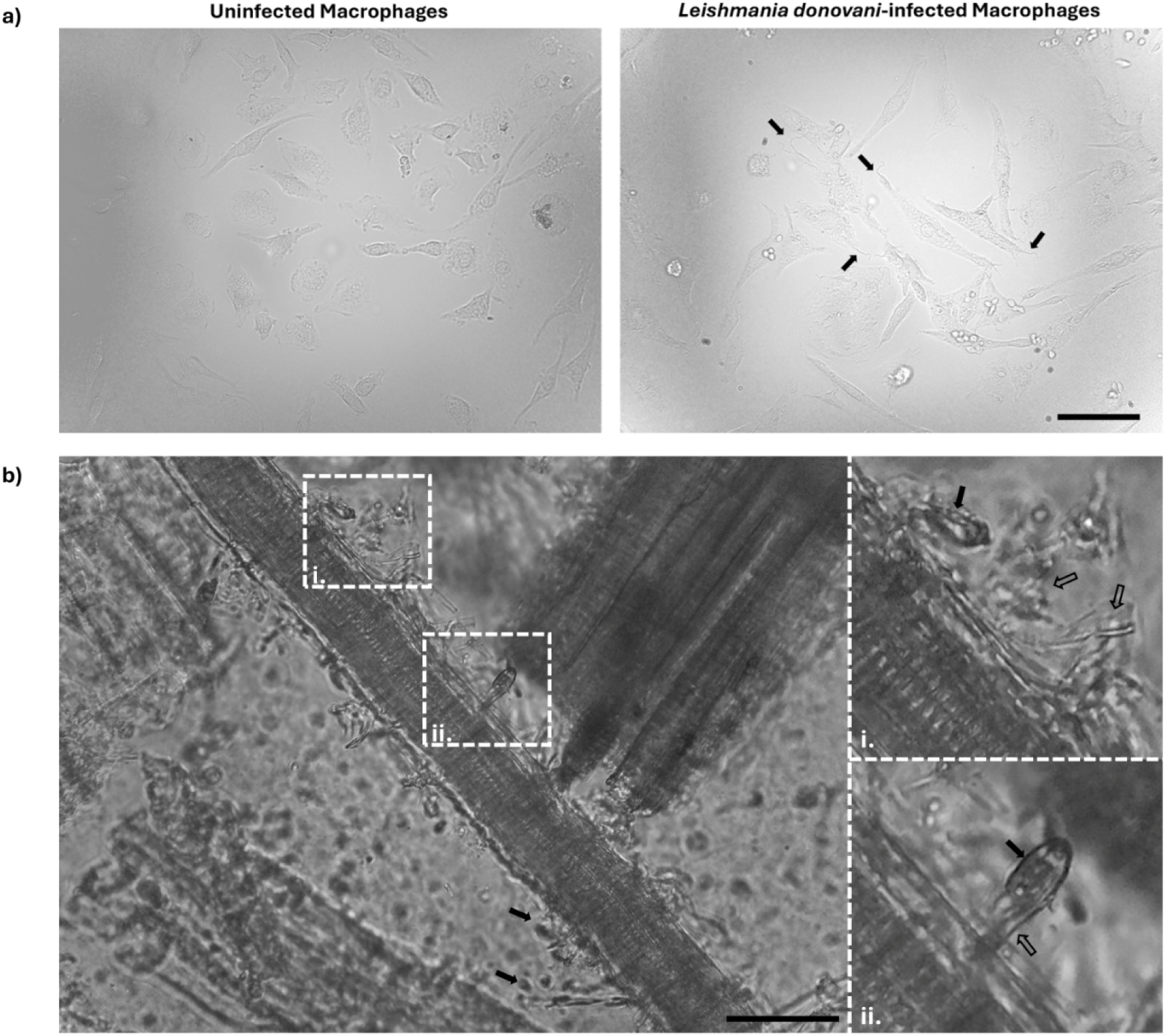
The LCI-OFM can be used to study diverse biological systems. (a) BMDMs were infected with promastigote forms of *L. donovani* or left uninfected. After 6 hours, non-internalised promastigotes were washed out and cultures were immediately returned to the incubator in a containment level 3 laboratory, before imaging with the LCI-OFM. Filled black arrows indicate flagella of promastigotes located outside macrophages, indicating ongoing parasite uptake. Scale bar = 50 µm. (b) Anaerobic gut fungi of the monocentric *Neocallimastix* genus were inoculated into a wheat-straw culture inside a vinyl anaerobic chamber and imaged after settling overnight. Hyphae are visible and have been labelled with open arrows, while the fungal head (sporangium) is clearly resolved using the LCI-OFM and labelled with filled black arrows. Scale bar = 50 µm.

We also assessed the LCI-OFM in a specialist anaerobic environment. Anaerobic gut fungi (AGF) are microorganisms typically found in the rumen of ruminant livestock and play an important role in the microbial communities of ruminants as they possess a diverse supply of enzymes to degrade lignocellulose ^28,29^. Uncharacterised anaerobic gut fungi of the monocentric *Neocallimastix* genus were cultured in wheat straw inside a type A vinyl anaerobic chamber (Coy Laboratory Products, MI, USA) and imaged after settling overnight. Despite the stringent anoxic conditions required for AGF, the LCI-OFM readily resolved key morphological features, including the sporangium and associated hyphae (Figure 6b), demonstrating its suitability for studying anaerobic species within complex environments.

Together, these examples demonstrate that the LCI-OFM delivers high-quality imaging across widely different biological contexts, highlighting its broad utility for addressing diverse biological questions by bringing the microscope directly to restrictive or specialised spaces.

## Discussion

Our LCI-OFM addresses important limitations of the original OFM ^18^, enabling its specific use as a LCI tool. To prevent corrosion of essential electronic components, we redesigned the 3D-printed instrument so that it can be safely housed and operated long-term in humid tissue culture incubators with the Raspberry Pi and Sangaboard no longer at risk of being exposed to unavoidable moisture. To ensure living cells can be maintained in optimal physiological conditions, we designed a simple stage adaption that can support 35 mm culture dishes that are necessary for proper perfusion of culture medium and supply of essential nutrients to cells. To further protect the health of living cells, we manufactured custom plugins containing current-limiting resistors that reduced thermal output of the stepper motors, preventing heat from building up in the enclosed incubators and preserving the delicate environmental conditions. To ensure rapid initiation of timelapse experiments, we created a simple timelapse-focused GUI that is intuitive, reliable and lightweight. Importantly, beyond installation of the LCI-GUI, no internet connection is required to operate this application. Critically, we have shown our LCI-OFM is capable of generating meaningful timelapse data, at the fraction of the cost of high-end systems. Our 3D-printed modifications are compatible with the original OFM, and do not affect its functionality. Therefore, users are able to use the original OFM for routine microscopy or incorporate our LCI adaptations to conduct timelapse experiments with ease, further enhancing the versatility of this 3D-printed tool. Our LCI-OFM further lowers the barrier to entry to advanced microscopy for researchers in LMICs who are disadvantaged when it comes to the adoption of modern imaging and image analysis techniques ^9–11^. The LCI-OFM could also provide opportunities for students pursuing life sciences in academic institutes globally, where studying living cells at a temporal resolution may be restricted due to financial constraints limiting access to dedicated timelapse hardware.

Opportunities to refine both the hardware and software components of the LCI-OFM remain. Printing in ASA, while offering substantial improvements to timelapse data quality, may represent a significant cost for researchers with tighter budgets. In settings where 3D-printing resources are available but material options are limited, lower cost filaments may be the only accessible options. Solutions to reinforce PLA could help mitigate the weakening of the material in incubators, including acrylic sprays to reduce moisture absorption, or increasing the wall thickness or infill during the print for structural or weight-bearing parts of the frame. Alternatively, printing with glycol-modified polyethylene terephthalate (PETG) may offer increased thermal stability with a similar cost as PLA. Future enhancements to the LCI-GUI could include autofocus routines before each new image acquisition, options for tiling and stitching to capture larger ROIs, or sampling multiple ROIs across a dish. We note that the OpenFlexure team is actively exploring timelapse acquisition capabilities within the core OFM software. While these efforts are ongoing and not yet fully validated or released, they do represent an important complementary route to expanding LCI functionality on this platform.

Lastly, we recognise the expansive efforts being made to democratise microscopy beyond the OFM project. Recent frugal and modular microscopy initiatives, from smartphone-based systems to fully customisable open-hardware platforms, highlight a global shift toward accessible, adaptable imaging technologies that broaden participation in research, teaching and diagnostics.

In conclusion, by redesigning the OFM for compatibility with humid, temperature-controlled incubators and for use in settings with limited resources, we have closed a critical gap between low-cost open-source microscopy and the practical demands of long-term LCI. Our hardware modifications to the OFM improve resilience to the harsh environment of standard tissue culture incubators and enhance mechanical stability, while the locally installed LCI-GUI enables fully offline, user-friendly rapid setup of timelapse experiments. Together, these modifications allow the OFM to generate biologically meaningful, analysis-ready datasets. This work expands the utility of the OFM for researchers in low-resource settings, supporting the generation of preliminary data, enabling deeper investigation of locally relevant biological questions, and strengthening pathways toward equitable access to advanced imaging technologies. We encourage the wider global microscopy community to adapt and build upon either the original OFM or the LCI-OFM, while maintaining open source practices so that all microscopists everywhere, regardless of resource, can benefit from continued technological advancement.

## Online Methods

### Breast Cancer Cell Line Culture

MCF-7 and MDA-MB-231 cell lines were used throughout the development of the LCI-OFM. Both breast cancer cell lines were maintained in Dulbecco’s Modified Essential Medium (DMEM; Gibco, S41966-029) supplemented with 5% (v/v) foetal bovine serum (FBS; Gibco, 10270106) in a humidified Binder CO_2_ incubator at 37 °C and 5 % CO_2_. MCF-7 cells were purchased from ATCC.

### Isolation and Differentiation of Murine Bone Marrow Macrophages

Bone marrow-derived macrophages (BMDM) isolated from BALB/c mice were differentiated in DMEM (Gibco, 11960044) supplemented with 20% hi-FCS (Gibco, A5256701), 1% L-glutamine (Gibco, 25030081), 20 ng mL -1 m-CSF (BioLegend, 576402), 1x MEM Non-Essential Amino Acids Solution (Gibco, 11140050), and 10 mM HEPES (Gibco, 15630080). BMDM cultures were maintained in DMEM medium supplemented with 10% hi-FBS and 10 mM L-glutamine (Gibco, 25030081) at 37 °C, in 5% CO_2_.

### Culture of *Leishmania donovani* Promastigotes

Promastigote forms of *Leishmania donovani* MHOM/ET/1967/HU3 line, also referred to as LV9 or HU3, were used to infect BMDMs. Promastigote parasites were cultured at 25 °C in M199 medium (Gibco, 12350039), supplemented with 10% heat-inactivated FBS (hi-FBS, Gibco, A5256701), 5 μg/mL hemin (Sigma-Aldrich, 51280-5G), 10 μM 6-biopterin (Sigma-Aldrich, B2517), and 100 U penicillin - 100 μg mL -1 streptomycin (Sigma-Aldrich, P4333), pH 7.2.

### Anaerobic Gut Fungi Culture

Monocentric *Neocallimastix* fungal cultures were maintained in sealed Hungate tubes using an adapted Medium C formulation ^30,31^ and subcultured every three days. Approximately 1 g of milled wheatstraw was added per 10 ml Hungate tube as the carbon source. Tubes were autoclaved prior to use, and chloramphenicol was added fresh before each transfer to a final concentration of 0.01%.

### Live-Cell Imaging of Breast Cancer Cells

For timelapse experiments on the LCI-OFM, breast cancer cells were seeded to a density of 3.2×10^4^ or 1×10^5^ cells per cellview glass-bottom 35 mm culture dish (Greiner, 627860). Cells were left to settle to the surface of the glass coverslip overnight before timelapse initiation. The LCI-GUI was used to set up standard timelapse experiments with a default interval of 6 min and a time course length of 24 or 48 h. For measuring thermal output of stepper motors, a Hikmicro Eco handheld superIR resolution camera was used.

### Docetaxel Treatment of MDA-MB-231 Cells

MDA-MB-231 cells were seeded to cellview dishes as described above. After cells had settled, culture medium was aspirated and cells were washed with 1X phosphate-buffered saline (PBS; Gibco, 14200-067) before replenishment of culture medium supplemented with either 0.01% DMSO (vehicle) or 1 μM docetaxel (Cambridge Bioscience, D5709). 48 hour timelapses with 6 min intervals were initiated using the LCI-GUI. Experiments were conducted for four biological repeats.

### Imaging of *Leismania-*Infected BMDMs in a Containment Level 3 Laboratory

For imaging *Leishmania*-infected and non-infected macrophages, 5×10^5^ differentiated BMDMs were seeded into 35 mm CellView glass-bottom culture dishes (Greiner, 627860). After 24 hours, macrophages were infected with late-logarithmic phase *Leishmania* promastigote at a multiplicity of infection (MOI) of 10 parasites per macrophage. Parasites were resuspended in DMEM supplemented with 5% heat-inactivated fetal calf serum (hi-FCS) prior to infection. Following 6 h of incubation at 37 °C and 5% CO₂, non-internalised promastigotes were removed by washing with the culture medium. Imaging was then initiated immediately after the washing step.

### Live-Cell Imaging of Monocentric *Neocallimastix*

Anaerobic *Neocallimastix* fungi were inoculated into µ-Slide 8-well chamber slides (Ibidi, 80806) in a straw medium supplemented with 0.1 ml of chloramphenicol (1% w/v stock; ACROS Organics, 56-75-7) per 10 ml culture medium, 0.6% w/v streptomycin sulphate (Apollo Scientific 3810-74-0) and 0.3% w/v penicillin V potassium salt (ACROS Organics 132-98-9). Inoculations were performed inside a type A vinyl anaerobic chamber (Coy Laboratory Products, MI, USA), with cultures incubated at 39°C and allowed to settle overnight before imaging with the LCI-OFM inside the anaerobic workstation.

### 3D-printing and modelling

All filament used for 3D-printing was purchased from bambulab.com. The Bambu Lab X1 and Bambu Lab P1S printers were used for printing out all 3D-printed parts with standard 0.2 mm PLA or ASA filament slicer settings. Autodesk Inventor Software was used for 3D modelling

### LCI-OFM Assembly and Electronics

Full details of the original OFM components, including assembly instructions can be found at build.openflexure.org. All 3D models for the LCI-OFM can be accessed, downloaded, modified and printed from Zenodo (DOI 10.5281/zenodo.17532866). Extension cables for the *x*, *y* and *z* stepper motors were assembled in-house using 2.5 mm connectors (JST, XH), including top-entry shrouded headers (JST, B5B-XH-A), high-box connector housing (JST, XHP-5) and crimp contact terminals (JST, SXH-001T-P0.6). Wiring consisted of 0.33 mm², 22 AWG, 7/0.25 mm PVC hook up wire (RS, 207-5303), cut to approximately 200 cm for both motor and LED extensions. New LED connectors were created using long break away headers (SparkFun, PRT-10158) and arduino stackable header kits (SparkFun, PRT-11417), each cut to two pin lengths. 200 cm Pi Camera ribbons were purchased from Pimoroni (Pimoroni, CAB0305). Custom plugins for the stepper motors were designed with four metal-film leaded resistors (Vishay, PR01) arranged in series with each winding of the stepper motor to limit the current draw while maintaining sufficient torque for motor movement. Heatshrink tubing and RS Pro 25 mm expandable braided PET black cable sleeve (RS, 408-271) were applied to exposed cabling.

### LCI-GUI Installation

Installation guidelines and full documentation for the LCI-GUI are available at https://doi.org/10.5281/zenodo.21379245.

### Cellpose

A custom cellpose model for image segmentation was trained using Cellpose v3.1.1.2 on brightfield images of MDA-MB-231 breast cancer cells captured on the LCI-OFM, and is available at https://github.com/uoy-research/cellphe-pipeline-images/blob/main/cellpose-3/models/CP_O FM_20260114

## Acknowledgments

We thank the OpenFlexure team for their constructive discussions regarding future timelapse capabilities for the OFM software. While these developments are ongoing and independent of the work presented here, our conversations helped us to better understand potential directions for expanding LCI functionality within the broader OpenFlexure community.

This work was supported by the Wellcome Trust (310891/Z/24/Z), GenerationResearch (N0017329/0002), EPSRC (EP/Z535114/1 and EP/S001581/1) and York Against Cancer.

## Supplementary Information

### Supplementary Files

Supplementary File 1 mods_to_openflexure

Supplementary File 2 ijmacro_openflexure_resize_8bit

### Supplementary Figures

**Supplementary Figure 1.**
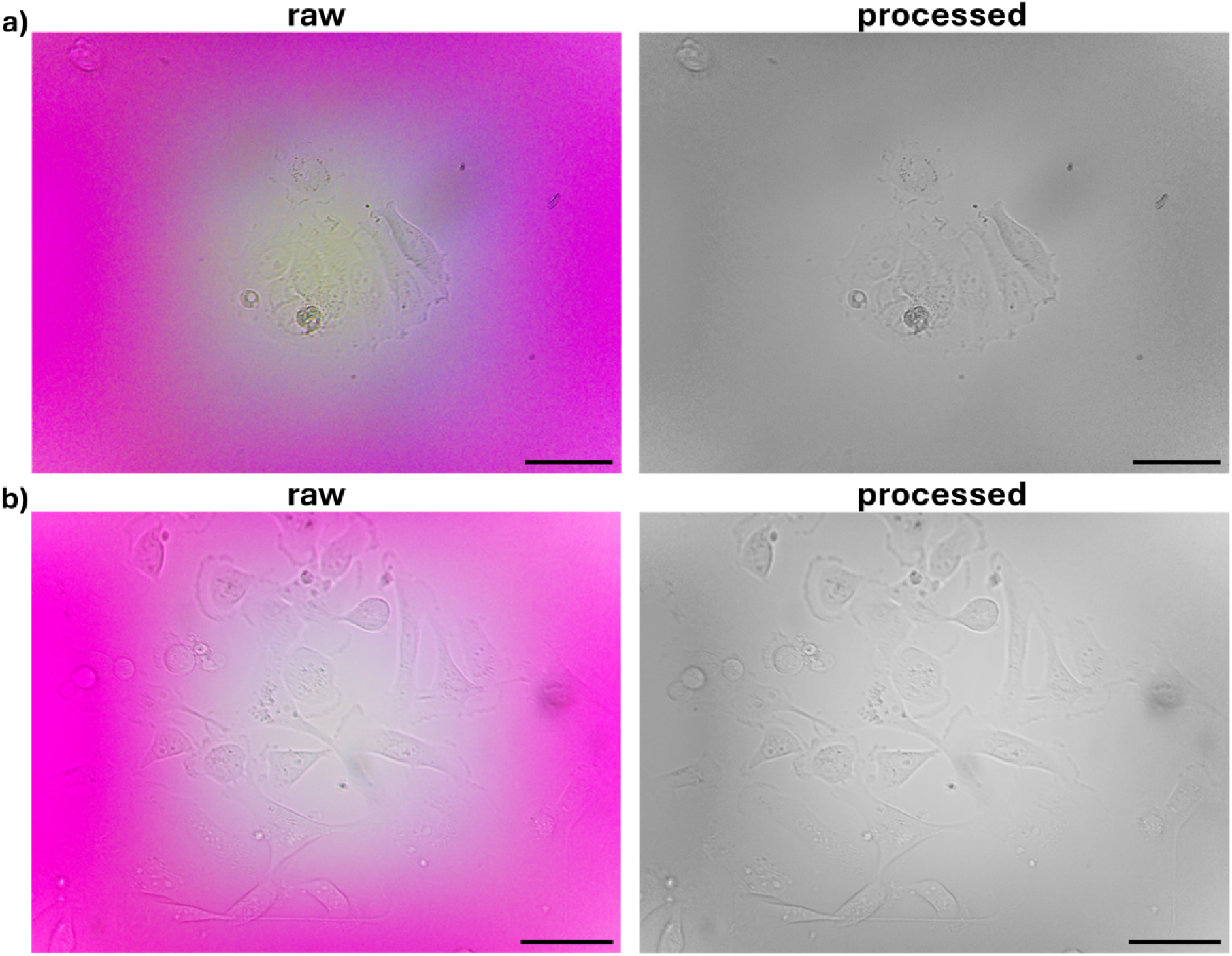
ImageJ macro to convert RGB to 8-bit greyscale and resize images post acquisition. Example images of a) MCF-7 and (b) MDA-MB-231 breast cancer cells captured using the LCI-OFM. Raw images are processed in imageJ using a custom macro to convert images into 8-bit greyscale and resize images to reduce the amount of storage required to maintain data. Scale bar = 50 µm.

**Supplementary Figure 2.**
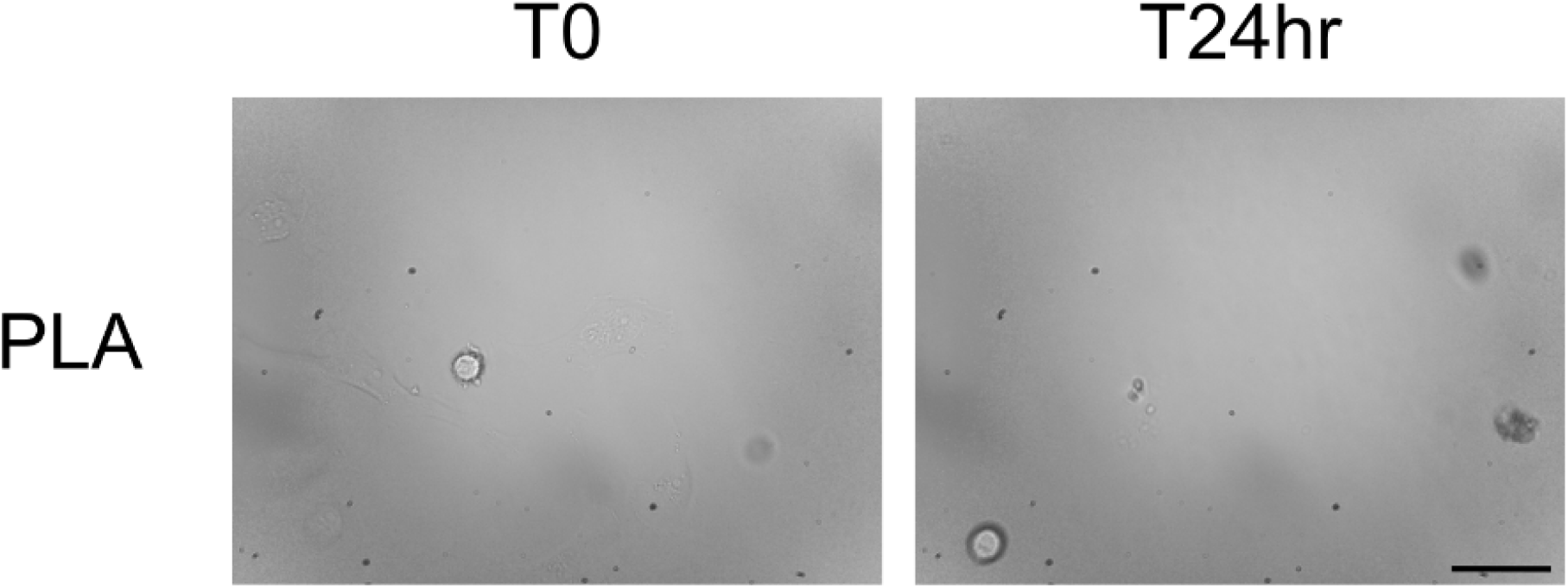
Drift in focus (PLA). MDA-MB-231 cells imaged for 24 h with 6 min intervals using a LCI-OFM printed in PLA. Scale bar = 50 μm.

**Supplementary Figure 3.**
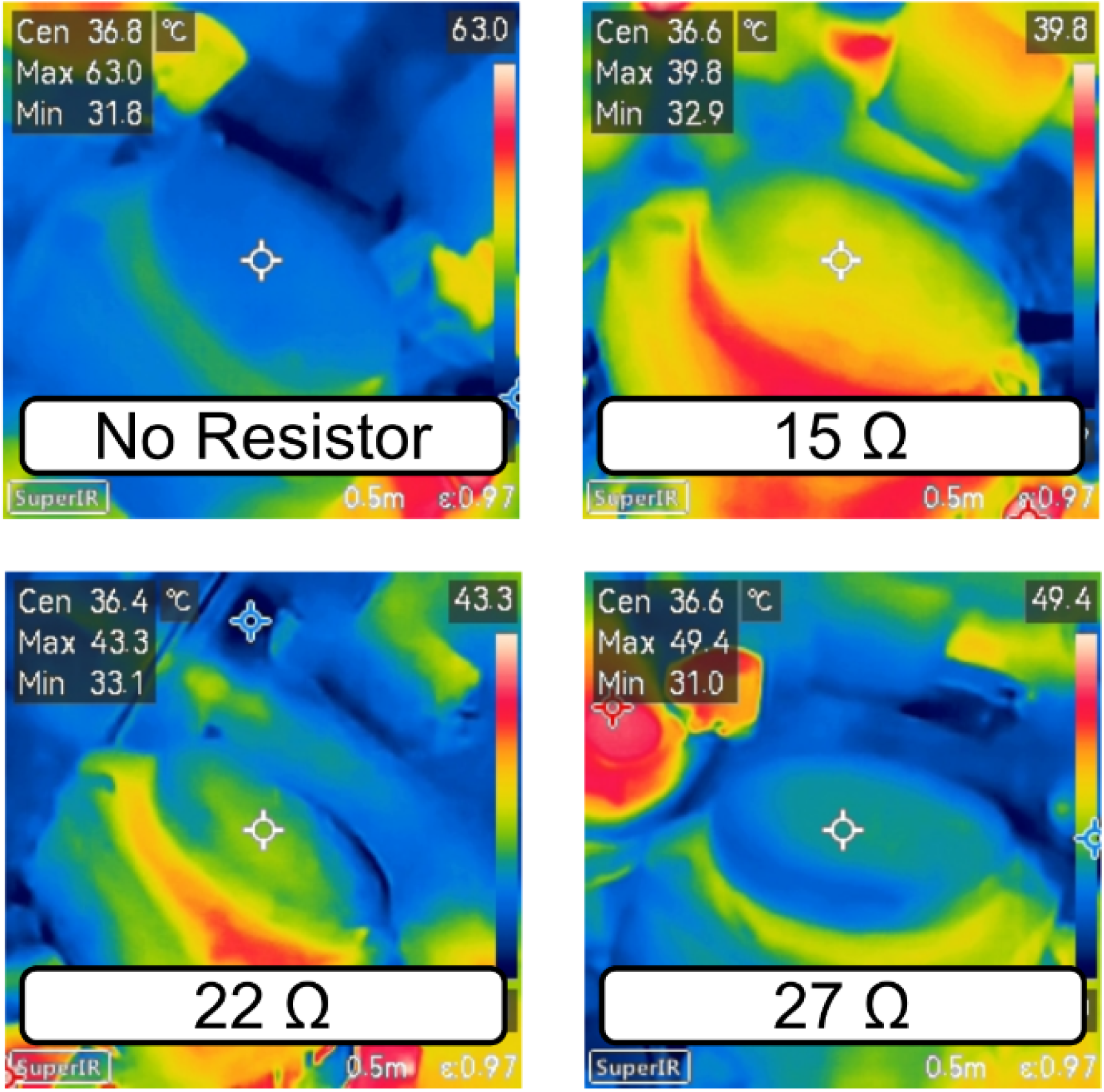
Final temperature of MDA-MB-231 cells in culture vessels at the end of a 48-hour timelapse experiment. MDA-MB-231 breast cancer cells were imaged on a LCI-OFM for 48 h with a 6 min interval. Stepper motor currents were either unperturbed, or were reduced with custom plugins containing either four 15 Ω, 22 Ω or 27 Ω resistors. The temperature of the cells and the culture vessel was captured at the end of the experiment and is shown by the “cen” value.

**Supplementary Figure 4.**
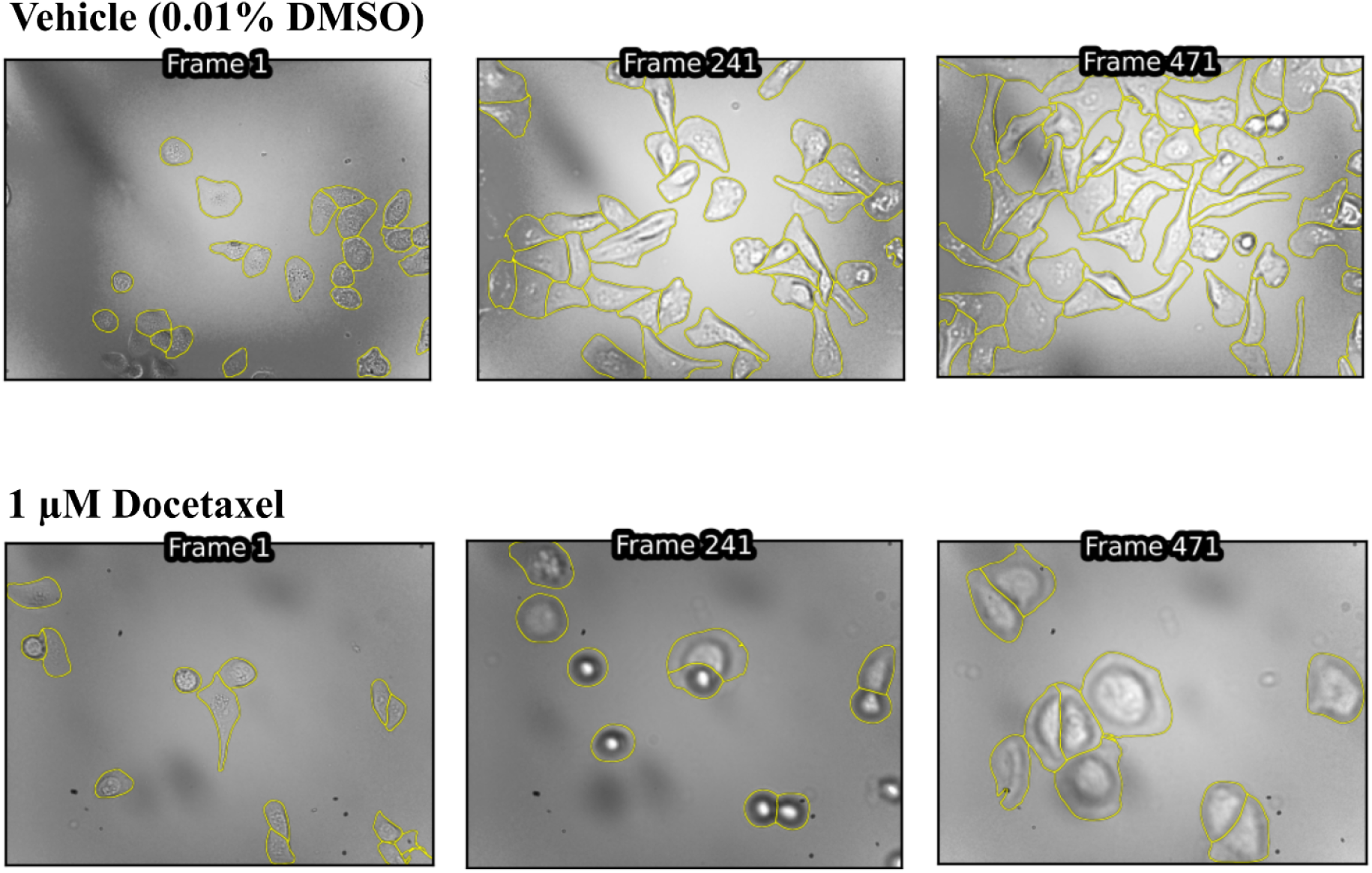
Example of segmentation masks of MDA-MB-231 cells treated with either vehicle or 1 μM docetaxel. Segmentation was performed using our custom Cellpose model.

### Supplementary Tables

**Supplementary Table 1.** Cost and material characteristics of common 3D-printing polymers evaluated for microscope use in 37°C humid tissue culture incubators. Characteristics are according to technical data sheets for each polymer, provided by bambulab.com. Cost reflects the price of each 1 kg spool available to purchase from bambulab.com at the time of writing. Abbreviations: Megapascals (MPa); Vicar Softening Temperature (VST); Great British Pound (GBP); International Organization for Standardization (ISO). Strength data are mean ± SD.

| Polymer | X-Y Tensile Strength (MPa); ISO 527, GB/T 1040 | X-Y Bending Strength (MPa); ISO 178, GB/T 9341 | VST (°C); ISO 306, GB/T 1633 | Cost (GBP/kg) |
| --- | --- | --- | --- | --- |
| PLA | 35 ± 4 | 76 ± 5 | 57 | 17.99 |
| PLA+ | 34.9 ± 1 | 65 ± 1 | 62 | 18.99 |
| ASA | 37 ± 3 | 65 ± 5 | 106 | 27.99 |
| PC-FR | 60 ± 4 | 90 ± 4 | 114 | 48.99 |

### Supplementary Videos

Supplementary Video 1 pla

Supplementary Video 2 pla_24hr

Supplementary Video 3 pla_tough_24hr

Supplementary Video 4 pcfr_24hr

Supplementary Video 5 asa_24hr

Supplementary Video 6 mdamb231_no_resistor

Supplementary Video 7 mdamb231_vehicle

Supplementary Video 8 mdamb231_docetaxel

